# Neural and behavioral adaptation to bilateral maps in primary somatosensory cortex

**DOI:** 10.1101/2025.06.10.658919

**Authors:** Varun B. Chokshi, Yi-Ting Chang, Daniel M. Ra, Reha S. Erzurumlu, Daniel H. O’Connor

## Abstract

Mice rely heavily on their sophisticated whisker somatosensory system to explore and navigate their surroundings. The primary whisker somatosensory cortex (wS1) receives contralateral sensory input due to a complete crossover of axonal projections ascending from the brainstem to the thalamus, resulting in a somatotopic map that exclusively represents the contralateral side of the face. This axonal crossover is disrupted in mice with a conditional knockout of the Robo3 gene, leading to abnormal bilateral representations of the whiskers in wS1. We explored the brain’s ability to adapt to a profound alteration of its somatotopic maps by using these Robo3 mutant mice. Performance on a discrimination task, in which mice reported whether a left-side or a right-side whisker was deflected, was on par with that of wild-type littermates. Unilateral optogenetic inhibition of wS1 showed that activity in the wS1 contralateral to a stimulated whisker was required for mice to report its side correctly, despite the representation of that whisker in the uninhibited hemisphere. Single-unit recordings in wS1 and the whisker primary motor cortex (wM1), a major downstream target of wS1, showed abnormal bilateral whisker responses in wS1 but largely normal responses in wM1, suggesting that the bilateral responses in wS1 were filtered out along the sensorimotor processing stream. Our results demonstrate that the brain can adapt to fundamental alterations in tactile input to construct accurate sensorimotor representations.

## Introduction

The whisker region of mouse primary somatosensory cortex (wS1) is a marquee model for relating developmental processes to cortical structure and function^1^. Each column of wS1 receives input from a unique whisker on the contralateral side of the face, in a sharply segregated area (barrel) of layer 4, via the primary thalamic ventral posterior medial (VPM) nucleus^2^. Inputs arise from the contralateral whiskers due to a complete crossover of projections from the brainstem principal trigeminal nucleus to VPM^3,4^. The result of this developmental process is a somatotopic map of the contralateral whiskers in layer 4 of wS1^5,6^.

In mice with a conditional mutation of the gene for the midline guidance receptor Robo3 in rhombomeres 3 and 5 (Krox20:cre;Robo^lox/lox^, Robo3cKO)^7^, this developmental process is disrupted and somatotopic maps in wS1 are no longer strictly contralateral^4^. Robo3cKO mice instead have bilateral whisker maps in layer 4 of wS1, receiving VPM inputs to a given barrel from either a contralateral or an ipsilateral whisker, without a change in the overall size of wS1^4^ (Figure 1A).

**Figure 1:**
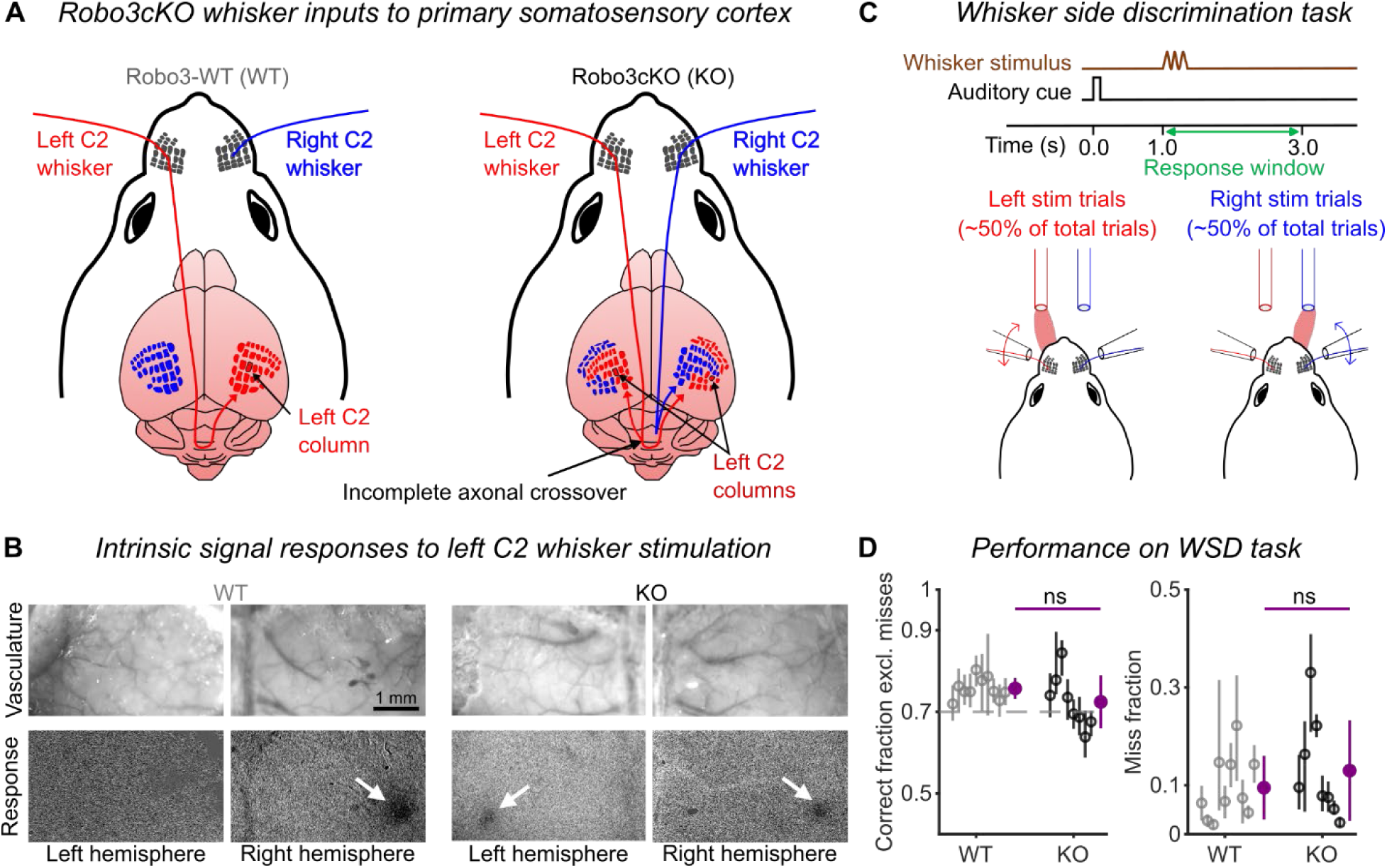
Robo3cKO mice perform a whisker side discrimination task. (A) Robo3cKO mice have bilateral whisker maps in each hemisphere. In WT mice (left), wS1 receives inputs from only contralateral whiskers. In KO mice (right), incomplete crossover of projections from brainstem to thalamus results in inputs from both ipsilateral and contralateral whiskers. Pathways are illustrated for a single whisker (C2). (B) Intrinsic signal imaging in wS1 shows responses (white arrows) to left C2 whisker stimulation only in the right hemisphere of WT mice (left panels), but in both hemispheres of Robo3cKO mice (right panels). (C) Whisker side discrimination (WSD) task. Each trial started with an auditory cue followed by a 20 Hz,150-ms sinusoidal whisker deflection applied to either the left or right C2 whisker. Mice reported the side of the whisker stimulus by licking at one of two lick ports during a 2-s response window. (D) Robo3cKO mice performed the WSD task as well as WT mice. Fraction of trials correct (left panel; excluding misses) and fraction of miss trials (right panel) were both similar for WT and Robo3cKO mice. Open circles: individual mice (mean ± 95% hierarchical bootstrap CI). Purple circles: means (± SD) for each genotype. Mann-Whitney U tests between genotypes show no significant (ns) differences. Gray dashed line: target performance level.

Beyond layer 4, sensory signals from a whisker propagate across cortical layers in wS1 and on to downstream areas to ultimately generate appropriate motor outputs^8,9^. One of the dominant outputs from wS1 contributing to this sensorimotor processing is the corticocortical pathway from wS1 to whisker primary motor cortex (wM1)^10,11^. In wild-type (WT) mice, deflections of a single (C2) whisker evoke a response in the contralateral wM1 that follows the initial wS1 response by a few (<10) milliseconds^12,13^, is blocked by inactivation of the C2 column in wS1^12^, and occurs at the site of axonal projections from the C2 column to wM1^12^. Feedforward drive from wS1 is thus a major determinant of wM1 activity.

Here, we investigated the impact of bilateral whisker maps on sensorimotor function in Robo3cKO mice. We tested the ability of Robo3cKO mice to differentiate between whisker stimuli delivered to either the left or the right side of the face. Surprisingly, these mice were able to report the side of a whisker stimulus as well as their WT littermates. Unilateral optogenetic silencing of wS1 showed that Robo3cKO mice, like WT mice, relied on activity in the wS1 contralateral to the stimulated whisker to determine its side. After these behavioral experiments, we measured the spiking responses of single units in both wS1 and wM1 of Robo3cKO and WT mice. Despite bilateral responses to a whisker deflection in wS1 of Robo3cKO mice, spiking activity was largely normal and contralateral in wM1, indicating that ipsilateral-whisker responses were filtered out during sensorimotor processing. The mouse brain can therefore adapt to dramatic alterations in the organization of tactile inputs.

## Results

We first confirmed that stimulation of a single whisker drives activity in both hemispheres of Robo3cKO mice using intrinsic signal imaging (Figure 1B). Stimulation of the left C2 whisker in Robo3cKO mice activated wS1 in both hemispheres (Figure 1B, right). In contrast, in WT littermates, the left C2 whisker only activated the right wS1 (Figure 1B, left). These whisker stimulus responses are consistent with the patterns of bilateral thalamocortical inputs in Robo3cKO mice^4^. Using the following behavioral, optogenetic silencing, and electrophysiological experiments, we tested whether and how Robo3cKO mice with bilateral whisker maps in wS1 can discriminate between stimulation of the same C2 whisker on opposite sides of the face.

### Bilateral whisker maps in wS1 do not impair whisker side discrimination

If deflection of a single whisker produces a bilateral response in wS1 of Robo3cKO mice, would these mice be able to discriminate between left and right whisker stimuli? To test this, we used a whisker side discrimination (WSD) task where, in each trial, we delivered a brief C2 whisker stimulus to either the left or the right side of the face in head-fixed mice (Figure 1C). Mice were trained to lick at one of two lick ports on each trial to indicate the side of the whisker stimulus (Figure 1C). Licks occurring during a 2-s response window starting with stimulus onset (Figure 1C) were used to calculate the fraction of trials in which the mice correctly indicated the side of the whisker deflection. Trials in which mice failed to lick either port were considered “miss” trials and analyzed separately. We titrated stimulus amplitudes to achieve a criterion performance range of 70-75% correct (Methods).

We hypothesized that Robo3cKO mice would be severely impaired in the WSD task. Surprisingly, we found no deficiency in their performance compared to their WT littermates (Figure 1D, left). There was no difference between the two genotypes in whisker stimulus amplitudes required to achieve criterion performance (Figure S1A). Nor was there a deficiency in the ability to detect the whisker stimulus per se (as opposed to attributing it to the correct side), as the missed trial proportions were similar between WT and Robo3cKO mice (Figure 1D, right). Robo3cKO mice did not exhibit biases to one side or the other, as the fractions of correct (Figure S1B) and missed (Figure S1C) trials were comparable when calculated separately for trials with left- or right-whisker stimulation.

Together, these results show that Robo3cKO mice are not impaired in detecting a whisker stimulus and assigning it to the correct side of the face.

### Activity in wS1 contralateral to a whisker stimulus is critical for side discrimination

Robo3cKO mice showed normal levels of performance in the WSD task (Figure 1D). In principle they could solve the task using activity from one or both hemispheres, or even subcortical structures. We therefore used an optogenetic strategy to test whether WSD task performance depended on wS1 activity in each hemisphere. Channelrhodopsin-2 (ChR2) was expressed bilaterally in the inhibitory neuronal population of wS1 by injecting an mDLX enhancer-driven viral expression system (rAAV-mDlx-ChR2-mCherry) in neonatal pups^14^ (Figure S2A). In each session, unilateral optogenetic inhibition of wS1 was performed in 20% of the trials (Figure 2A) using a clear skull preparation^15^ (one hemisphere per session; Methods). Optogenetic inhibition spanned from 200 ms prior to whisker stimulus onset through the end of the response window (Figure 2A).

**Figure 2:**
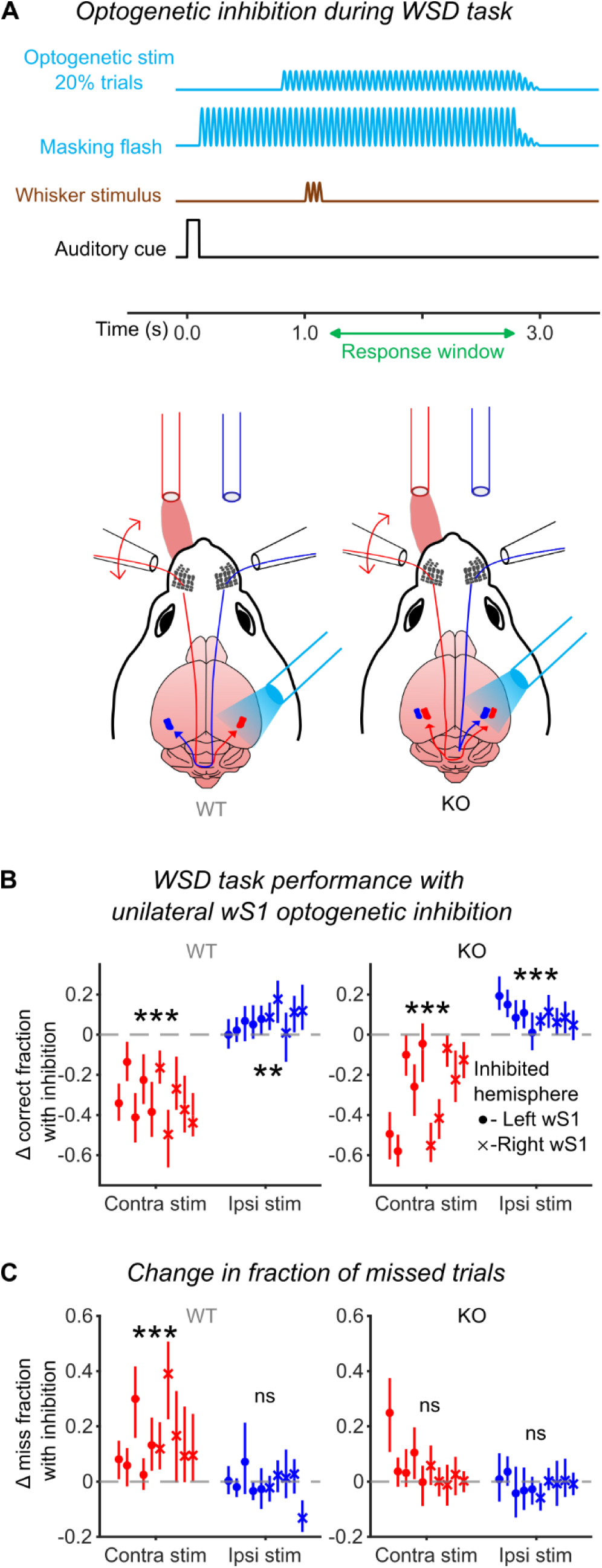
Contralateral wS1 activity is critical for reporting the side of a whisker stimulus. (A) Optogenetic inhibition experiment. The left or right hemisphere was silenced in 20% of trials by placing an optic fiber over wS1 to excite inhibitory neurons. (B) Performance on trials with whisker stimuli contralateral to the inhibited hemisphere (red symbols) was reduced for both WT (left panel) and Robo3cKO (right panel) mice. A slight improvement occurred on trials with whisker stimuli ipsilateral to the inhibited hemisphere (blue symbols), for both genotypes. (C) WT but not Robo3cKO mice miss more on trials with whisker stimuli contralateral to the inhibited hemisphere. (B-C) Each data point depicts a silencing experiment for one hemisphere (mean ± 95% hierarchical bootstrap CI). Circles: left hemispheres. Crosses: right hemispheres. Asterisks indicate significant binomial test results. ***, p < 0.001; **, p < 0.01; ns, p ≥ 0.05.

For WT mice, WSD task performance degraded on trials with whisker stimuli contralateral to the inhibited hemisphere (Figure 2B, left), to chance levels (Figure S2B). This was associated with an increase in missed trials (Figure 2C, left). Additionally, performance on trials with whisker stimulation ipsilateral to the inhibited hemisphere marginally improved in WT mice (Figure 2B, left). Together, these results in WT mice indicate that activity in the hemisphere contralateral to the stimulated whisker is critical for stimulus detection and assignment to the correct side. Surprisingly, unilateral optogenetic inhibition of wS1 in Robo3cKO mice produced a similar phenotype as in WT mice (Figure 2B). Inhibition degraded performance for stimuli contralateral to the inhibited hemisphere (Figure 2B, right), to chance levels (Figure S2C). There was however no consistent increase in miss trials (Figure 2C, right), in contrast to WT (Robo3cKO vs WT unpaired t-test, p = 0.03, comparison of red points in Figure 2C). This result suggests that ectopic activity in the uninhibited hemisphere ipsilateral to the whisker stimulus might be adequate to allow the Robo3cKO mice to detect a stimulus but not to assign its side correctly. Performance on trials with whisker stimulation ipsilateral to the inhibited hemisphere marginally improved in Robo3cKO mice (Figure 2B, right), as in the WT mice (Figure 2B, left). The amplitudes of the whisker stimuli on each side were similar for both genotypes for inhibition sessions (Figure S2D).

Together, these results show that, in both Robo3cKO and WT mice, activity in the hemisphere contralateral to a stimulus is critical for its assignment to the correct side.

### Segregated ipsilateral and contralateral whisker selectivity in wS1

Robo3cKO mice were remarkably like WTs in their ability to solve the WSD task and in the behavioral impact of wS1 inhibition. One explanation for these results could be that spiking responses across the population of wS1 neurons in trained Robo3cKO mice were also unexpectedly normal. This cannot be ruled out from prior studies with Robo3cKO mice because these relied on layer 2/3 calcium imaging^16^ and voltage-sensitive dye imaging^17,18^, which both relate indirectly to spiking, and did not monitor activity across all cortical layers. Mice in these prior studies were also not trained in the WSD task.

We made single-unit recordings using laminar silicon probes to measure spiking responses to whisker stimuli throughout the layers of wS1 in our trained mice (Figure 3A). We performed these recordings under light anesthesia to prevent possible contamination of deflection responses by self-generated whisker motions. We targeted recordings to both ipsilaterally and contralaterally responsive wS1 columns in Robo3cKO mice, using coordinates obtained from intrinsic signal imaging (Figure 1B). On each trial, either the left or right C2 whisker was stimulated with one of nine stimulus types chosen randomly. Left- and right-side stimulations were alternated across trials. The nine stimulus types comprised 1, 2 or 3 cycles of a sinusoidal waveform at 10, 20, or 40 Hz. This stimulus set included the WSD task stimulus (3 cycles of a 20 Hz sinusoidal deflection). For subsequent analyses, we included only units with significant responses occurring during this WSD task stimulus (i.e. to the 20 Hz stimulus occurring within a 150-ms window beginning with stimulus onset).

**Figure 3:**
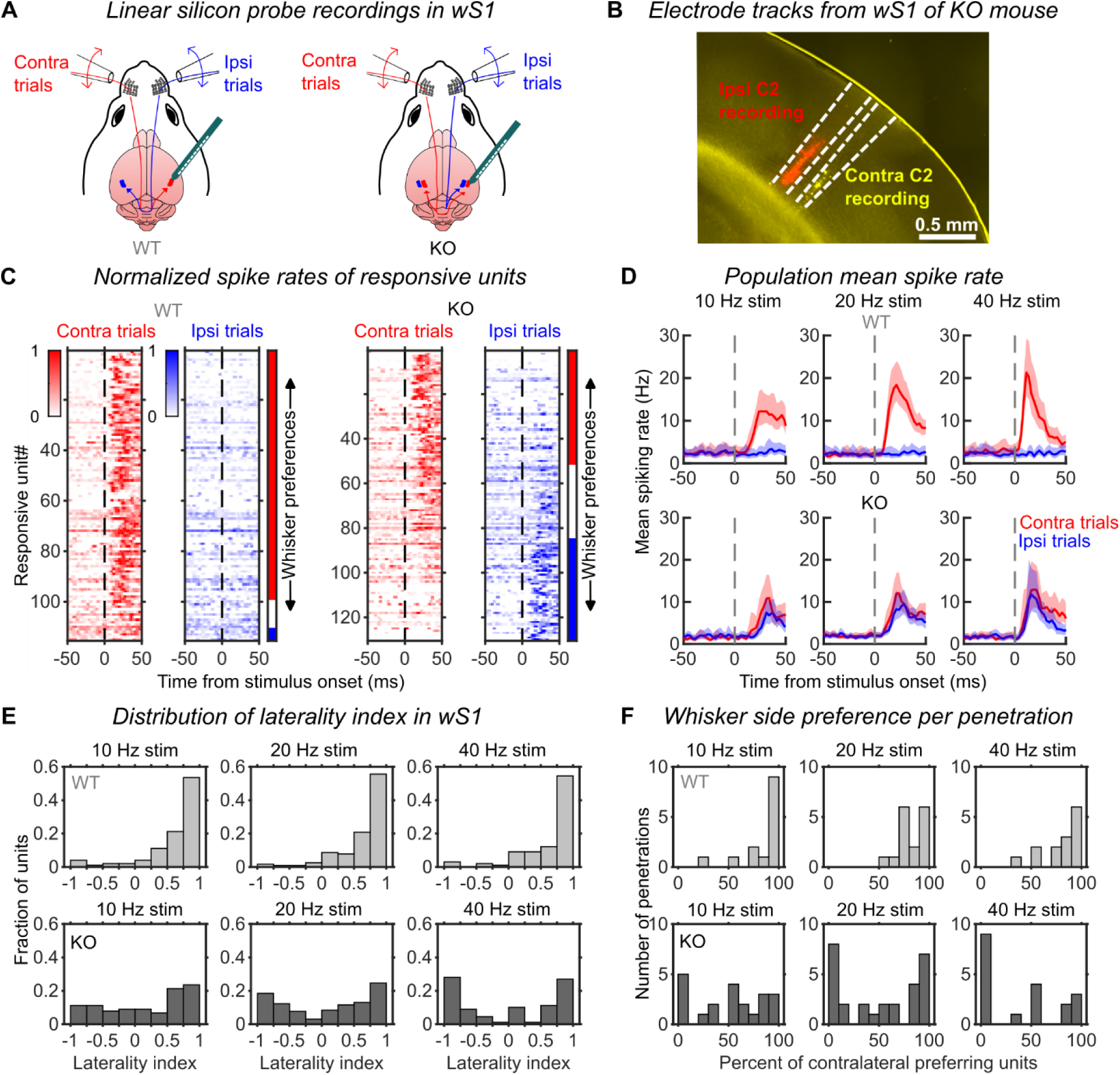
Ipsilateral and contralateral whisker responses in wS1. (A) Silicon probes were used in wS1 to record single-unit responses to stimulation of either the left or right C2 whisker. (B) Example histological section showing contralaterally and ipsilaterally responsive probe penetration sites in wS1 of a Robo3cKO mouse. Scale bar: 0.5 mm. (C) Normalized mean responses of wS1 units to all contralateral (red) and ipsilateral (blue) stimuli (10, 20 and 40 Hz frequencies and 1, 2 and 3 cycles). Units are sorted by whisker preference based on laterality index values. Vertical bars at right indicate units with contralateral (red), bilateral (white) or ipsilateral (blue) preferences. (D) Mean spike rate (± 95% CI) for contralateral (red) and ipsilateral (blue) whisker stimuli in WT (top row) and Robo3cKO (bottom row) mice, and for three frequencies of stimulation (columns). (E) Laterality index values from WT mice (top row) are unimodally distributed and highly positive, indicating a consistent preference for the contralateral C2 whisker. Values from Robo3cKO mice (bottom row) are more bimodally distributed with both negative and positive values, indicating preferences for either the ipsilateral or contralateral C2 whisker. (F) Percent of units in each recording penetration that preferred contralateral C2 stimulation for WT (top row) and Robo3cKO (bottom row) mice.

In WT mice, as expected almost all wS1 units responded more strongly to the contralateral rather than ipsilateral C2 whisker stimulus (Figure 3C, left). In contrast, wS1 units recorded in Robo3cKO mice responded either to the contralateral or to the ipsilateral whisker stimulus (Figure 3C, right), and this was true across stimulus frequencies and cortical depth (Figures S3-S4). This finding is consistent with anatomical projections from VPM^4^ and previous layer 2/3 calcium imaging results from Robo3cKO mice^16^. Most units were responsive to only one of the two whisker stimulus sides (96/128 units), although a smaller fraction was bilaterally responsive (32/128 units). Mean evoked responses increased with stimulus frequency for contralateral whisker stimuli in WTs (Figure 3D, top row, 10 Hz vs 40 Hz, paired t-test p = 0.005), and for both contralateral (10 Hz vs 40 Hz, paired t-test p = 0.00004) and ipsilateral whisker stimuli (10 Hz vs 40 Hz, paired t-test p = 0.00014) in Robo3cKOs (Figure 3D, bottom row). In Robo3cKO mice, the mean evoked responses of the wS1 population for ipsilateral and contralateral 40 Hz whisker stimuli were similar (Figure 3D, bottom row).

Thus, the spiking activity in wS1 of Robo3cKO is driven by whiskers on both sides of the face, with similar maximum evoked rates for contralateral and ipsilateral stimuli.

Anatomical projections in Robo3cKO mice from contralateral and ipsilateral whiskers are segregated into different barrels within cortical layer 4^4^. To quantify the degree to which responses to contralateral and ipsilateral whisker stimuli were segregated among the populations of recorded neurons, we calculated a “laterality index” (LI) for each unit. Positive LI values indicate a contralaterally selective response, and negative LI values indicate ipsilateral selectivity. In WT mice, LI values were distributed unimodally with over half the values >0.75 across all stimulation frequencies, indicating strong contralateral selectivity (Figure 3E, top). In contrast, in Robo3cKO mice, LI values were bimodally distributed with over half the values located either < -0.75 or > 0.75, indicating that individual units tended to be either strongly contralaterally selective or strongly ipsilaterally selective (Figure 3E, bottom). Distributions of LI indicated more ipsilaterally selective units across cortical depths (Figure S4B).

These analyses of LI show that individual single units were highly selective for either the contralateral or ipsilateral whisker. To assess whether neurons with ipsilateral or contralateral selectivity were organized into columns, we calculated for each recording penetration (made approximately perpendicular to the cortical surface) the percent of units with contralateral preference. Penetrations tended to have high percentages of contralaterally preferring units in WT mice (Figure 3F, top) and more bimodally distributed percentages in Robo3KO mice (Figure 3F, bottom). These results are consistent with a columnar organization of whisker preference. Together these results indicate that individual neurons in Robo3KO mice tend to have strong selectivity for either contralateral or ipsilateral whisker stimulation and that neurons with similar selectivity are segregated into different columns.

### Single-neuron and population-level discrimination of whisker side in wS1

Activity in wS1 was critical for performance of our WSD task in both WT and Robo3cKO mice (Figure 2). This suggests that neuronal activity in wS1 could be used to discriminate the side of C2 whisker stimulation on a trial-by-trial basis. To test this possibility, we performed receiver operating characteristic (ROC) analysis on the single-unit spiking of individual neurons in successive 50-ms time bins. We used trials with 20 Hz stimulation, as this was the stimulus frequency used in the WSD task. Prior to stimulus onset, as expected no units could discriminate between trials with ipsi- vs contralateral C2 whisker stimulation in either WT or Robo3cKO mice (Figure 4A, left panels; Figure S5A,B, left panels). In the 50-ms window starting with stimulus onset, however, 54.8% of units in WTs had single-trial activity that could discriminate significantly above chance (95% CI on AUC not including 0.5), with all these units selective for the contralateral C2 (AUC > 0.5; Figure 4A, middle top; Figure S5A, middle). In Robo3cKO mice, ∼35% of units could discriminate above chance in this same window, with similar numbers selective for the ipsilateral (AUC < 0.5; 16.9%) and contralateral (18.5%) C2 whiskers (Figure 4A, middle bottom; Figure S5B, middle). The fractions of units selective for ipsilateral and contralateral C2 whiskers differed significantly between genotypes (Figure S5C).

**Figure 4:**
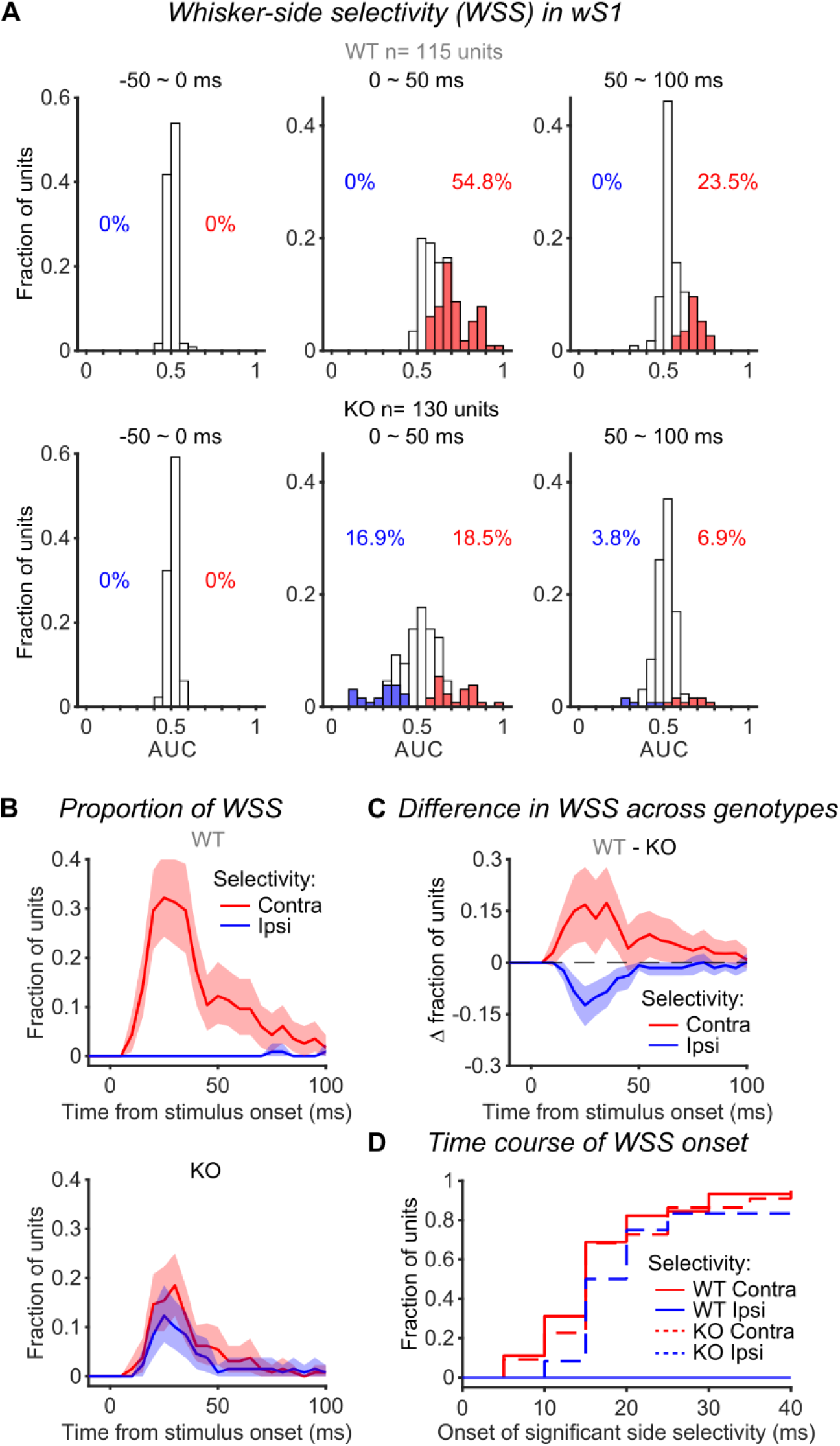
Trial-by-trial discrimination of whisker side by single neurons in wS1. (A) Area under the ROC curve (AUC) values quantifying how well the single-trial activity of each wS1 unit can be used to discriminate contralateral vs ipsilateral C2 whisker stimulation. Units with values significantly above (contralateral selective, red bars) and below (ipsilateral selective, blue bars) chance level (= 0.5) are indicated, along with the percentages of these units (red and blue text). Separate plots show results for different 50-ms time windows relative to stimulus onset (columns) and for WT (top row) and Robo3cKO (bottom row) mice. (B) Proportions of wS1 units showing contralateral (red) or ipsilateral (blue) selectivity at each time bin relative to stimulus onset, for WT (top) and Robo3cKO (bottom) mice. Error shading: 95% bootstrap CI (C) Difference between WT and Robo3cKO mice for fractions of units showing contralateral and ipsilateral selectivity at each 5-ms time bin. Error shading: 95% bootstrap CI. (D) Times relative to stimulus onset at which each wS1 unit first showed significant selectivity for the contralateral or ipsilateral C2 whisker. Numbers of selective units: WT contra = 45, WT ipsi = 0, KO contra = 22, KO ipsi = 12. Two-sided Kolmogorov-Smirnov tests: WT contra vs KO contra: p = 0.999; KO contra vs KO ipsi, p = 0.936; WT contra vs KO ipsi, p = 0.650.

We next repeated the ROC analyses but in shorter, 5-ms time bins, and calculated the fractions of units in each bin showing significant selectivity for the ipsilateral and contralateral C2 whiskers (Figure 4B), and the differences in these fractions across genotypes (Figure 4C). Development of selectivity across the population of single units followed a similar time course for units selective to each whisker and in both genotypes (Figure 4B). Robo3cKO mice, however, showed a lower fraction of units selective for the contralateral C2, because the total fraction of selective units was split between those with contralateral and ipsilateral selectivity (Figure 4B,C). Analysis of the time bins at which each unit first showed significant selectivity (Methods) was similar across the populations of units selective for ipsi- and contralateral C2 (Figure 4D). Together, these results show that the trial-by-trial activity of wS1 neurons in Robo3cKO mice can be used to discriminate between left- and right-side C2 whisker stimulation, with information about each side becoming available equally rapidly and with a time course similar to that of WT mice.

Behavioral discrimination of the side of whisker stimulation is likely to rely on the readout of activity across the population of neurons in wS1. To determine how well population activity could be read out from wS1 of Robo3cKO mice to discriminate between ipsi- and contralateral C2 whisker stimulation, we used linear discriminant analysis (LDA). We used LDA to classify trials into ipsi- vs contralateral stimulation trials based on the activity of simultaneously recorded single units in 50-ms time bins (Figure 5A). Prior to stimulus onset, as expected, populations could not be used to discriminate between trials with ipsi- vs contralateral C2 whisker stimulation in either WT or Robo3cKO mice (Figure 5A, left panels). In the 50-ms window starting with stimulus onset, however, population activity allowed classification of trials above chance in both genotypes, with median accuracies across sessions of 0.72 and 0.80 for Robo3cKO and WT mice, respectively (Figure 5A, right panels). We calculated 95% confidence intervals on the average classification accuracies across sessions and found that these overlapped for Robo3cKO and WT mice ([0.62, 0.76] vs [0.70, 0.81], respectively; Figure 5B).

**Figure 5:**
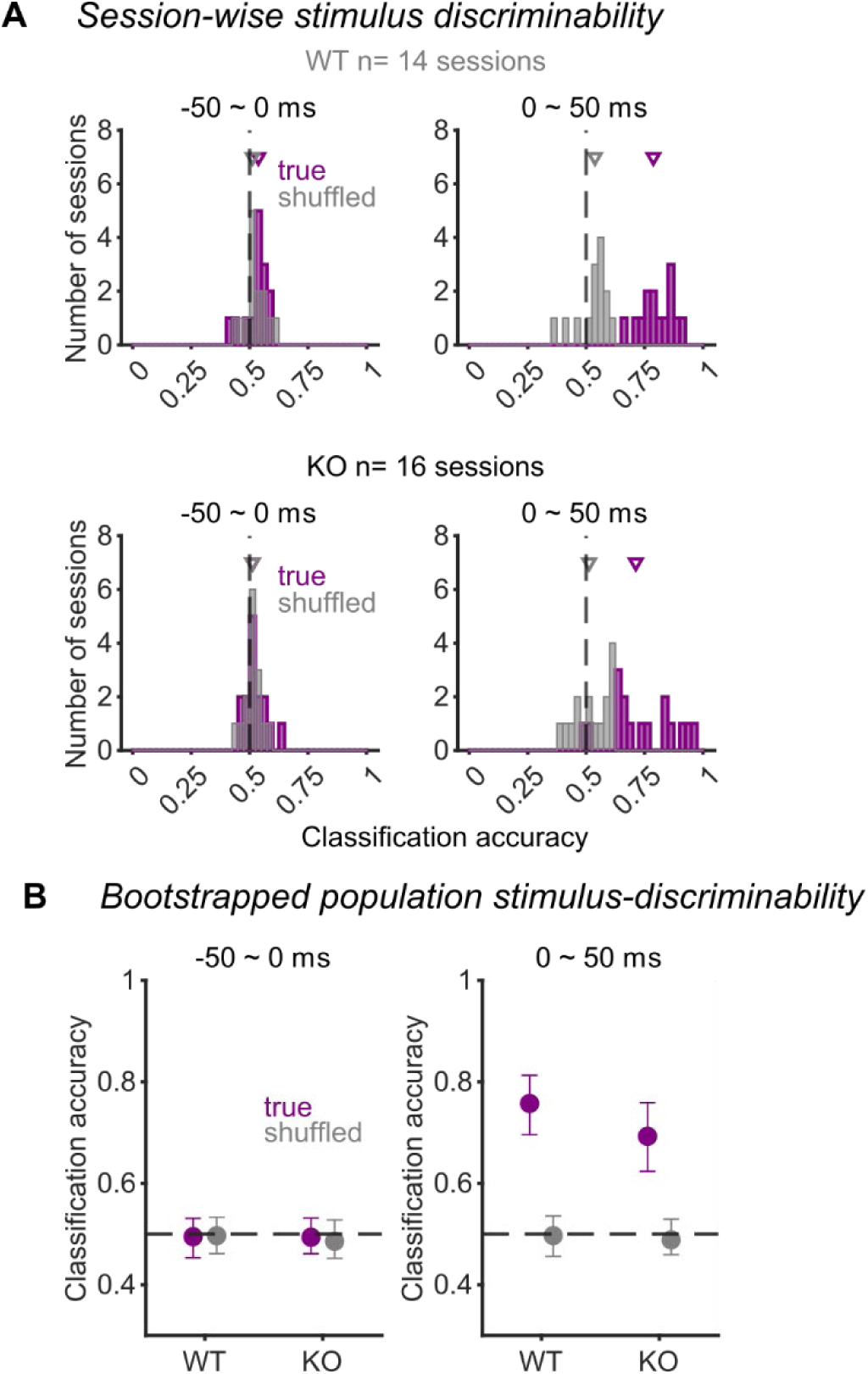
Population-level discrimination of whisker side by wS1. (A) Linear discriminant analysis (LDA) classification accuracies for contra- vs ipsilateral whisker stimuli for true (purple) and shuffled (gray) data. Separate plots show results from 50-ms analysis windows preceding (left column) or following (right column) onset of the stimulus and for WT (top row) and Robo3cKO (bottom row) mice. Arrowheads indicate medians. (B) Classification accuracies across sessions (bootstrapped mean ± 95% CI). Conventions as in panel A.

Together, these results show that despite the bilateral wS1 responses to single whisker stimulation in Robo3cKO mice, a simple linear decoding of the population activity could achieve performance levels similar to those of WT mice.

### Normal contralaterally selective responses in wM1

Population activity in wS1 could be linearly decoded to classify the side of a C2 whisker stimulus as well in Robo3cKO mice as in WT mice (Figure 5). This suggests that a downstream brain area could read out wS1 activity from Robo3cKO mice in a manner that separates contralateral and ipsilateral whisker responses. We tested this possibility by recording single-unit activity in wM1 using four-shank Neuropixel 2.0 probes^19^ targeted stereotactically (Figure S6). In WT mice, most units responded to the contralateral stimulus, with a minority of units responsive to the ipsilateral stimulus (Figure 6A, left). Whisker responses in wM1 of Robo3cKO mice were similar to WT mice, with most units being contralaterally responsive (Figure 6A, right). Mean responses across the population were larger for the contralateral compared with the ipsilateral C2 whisker across stimulation frequencies (Figure 6B). We repeated the ROC analyses that we previously performed for wS1. These showed that the trial-by-trial activity of 9.5% and 14.8% of single units could be used to discriminate ipsi- vs contralateral stimulation trials in the first 50 ms after stimulus onset in the Robo3cKO and WT mice, respectively (Figure 6C; Figure S8). Selectivity for the contralateral C2 whisker developed earlier and to higher levels than selectivity for the ipsilateral whisker, similarly for both Robo3cKO and WT mice (Figure 6D-F).

**Figure 6:**
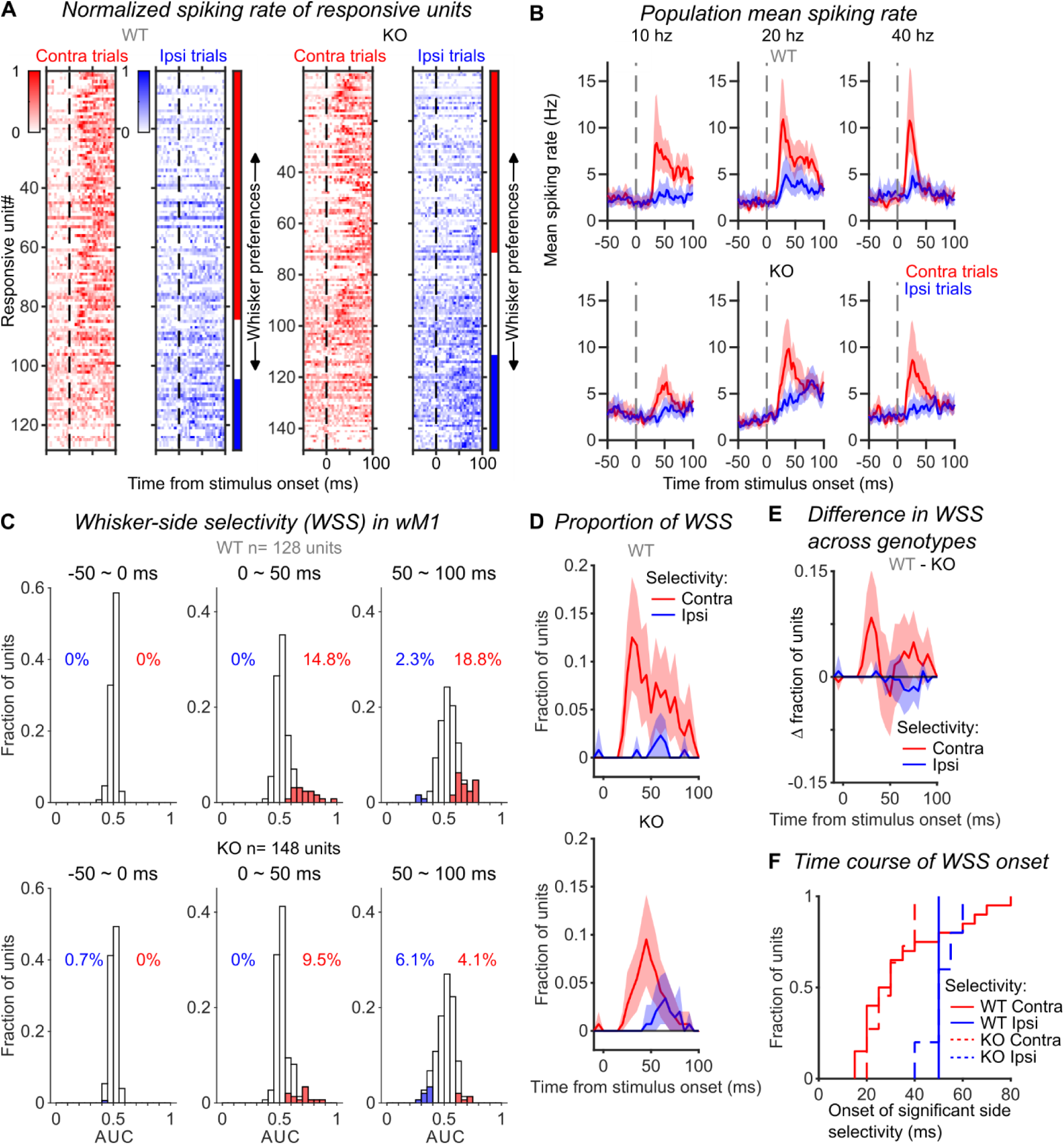
Contralaterally selective whisker responses in wM1. (A) Normalized mean response of wM1 units to all contralateral (red) and ipsilateral (blue) stimuli (10, 20 and 40 Hz frequencies and 1, 2 and 3 cycles). Units are sorted by whisker preference based on laterality index values. Vertical bars at right indicate units with contralateral (red), bilateral (white) or ipsilateral (blue) preferences. (B) Mean spike rate (± 95% CI) for contralateral (red) and ipsilateral (blue) whisker stimuli in WT (top row) and Robo3cKO (bottom row) mice, and for three frequencies of stimulation (columns). (C) AUC values quantifying how well the single-trial activity of each wM1 unit can be used to discriminate contralateral vs ipsilateral C2 whisker stimulation. Units with values significantly above (contralateral selective, red bars) and below (ipsilateral selective, blue bars) chance level (= 0.5) are indicated, along with the percentages of these units (red and blue text). Separate plots show results for different 50-ms time windows relative to stimulus onset (columns) and for WT (top row) and Robo3cKO (bottom row) mice. (D) Proportions of wM1 units showing contralateral (red) or ipsilateral (blue) selectivity at each time bin relative to stimulus onset, for WT (top) and Robo3cKO (bottom) mice. Error shading: 95% bootstrap CI. (E) Difference between WT and Robo3cKO mice for fractions of units showing contralateral and ipsilateral selectivity at each 5-ms time bin. Error shading: 95% confidence interval. (F) Times relative to stimulus onset at which each wM1 unit first showed significant selectivity for the contralateral or ipsilateral C2 whisker. Numbers of selective units: WT contra = 20, WT ipsi = 1, KO contra = 11, KO ipsi = 5. Two-sided Kolmogorov-Smirnov test of WT contra vs KO contra: p = 0.701. One-sided Kolmogorov-Smirnov tests: KO contra < KO ipsi: p = 0.005; WT contra < KO ipsi: p = 0.010.

Population activity in wM1 could be used to classify trials into those with ipsi- vs contralateral C2 whisker stimulation, with median classification accuracies across sessions of 0.64 and 0.76 in Robo3cKO and WT mice, respectively (Figure 7A; 100-ms time bins). Confidence intervals on the average classification accuracies across sessions overlapped for Robo3cKO ([0.55, 0.68]) and WT ([0.61, 0.77]) mice (Figure 7B).

**Figure 7:**
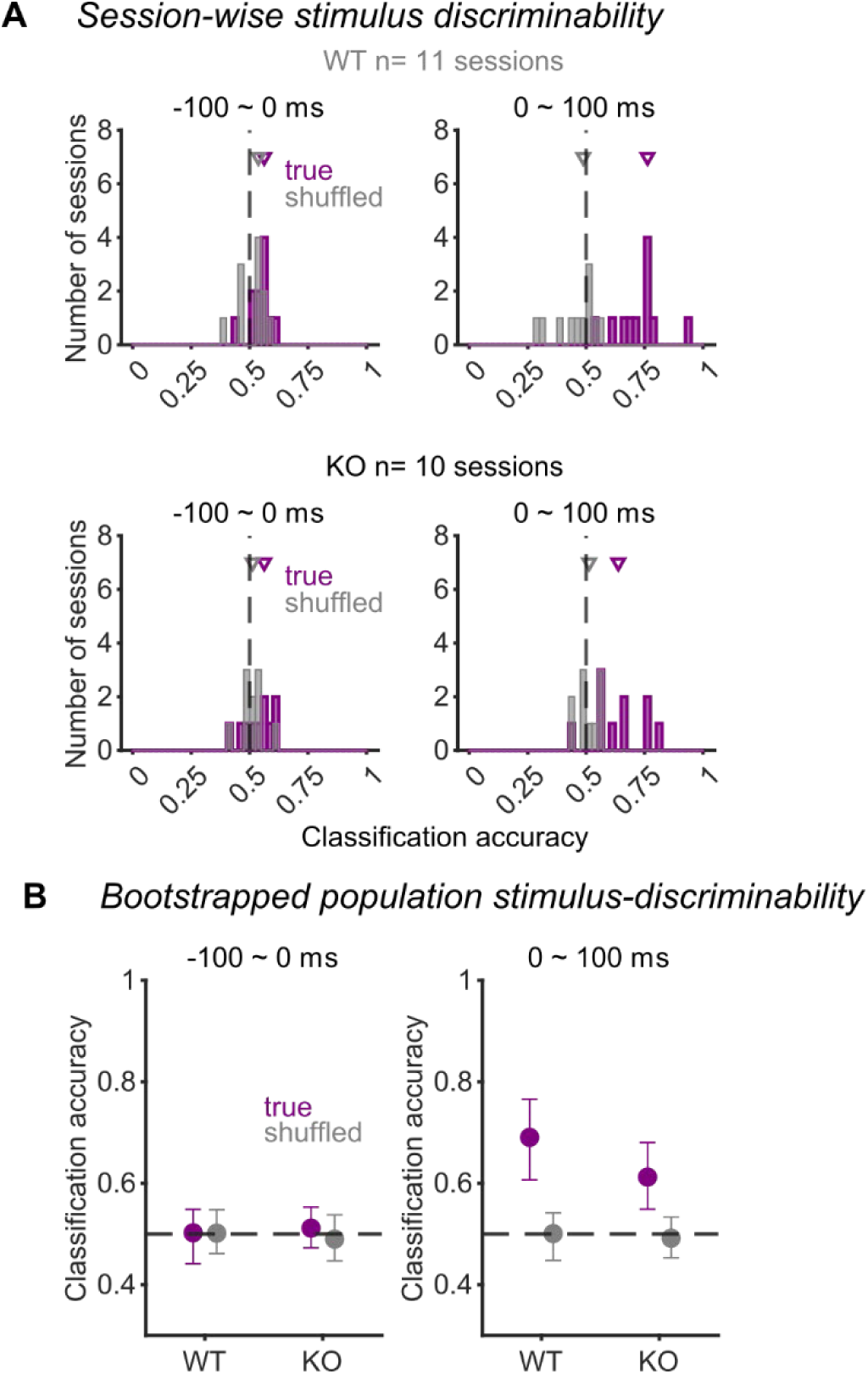
Population-level discrimination of whisker side by wM1. (A) LDA classification accuracies for contra- vs ipsilateral whisker stimuli for true (purple) and shuffled (gray) data. Separate plots show results from 100-ms analysis windows preceding (left column) or following (right column) onset of the stimulus and for WT (top row) and Robo3cKO (bottom row) mice. Arrowheads indicate medians. (B) Classification accuracies across sessions (bootstrapped mean ± 95% CI). Conventions as in panel A.

Together these data show that spiking responses in wM1 of Robo3cKO mice are largely contralaterally selective, as in WT mice. This contrasts with the dramatic differences between Robo3cKO and WT mice observed in wS1.

## Discussion

In contrast to senses such as vision and audition, ascending sensory pathways for touch remain fully lateralized through the level of primary sensory cortex. In Robo3cKO mice, however, prior work has shown that stimulation of a single C2 whisker evokes responses in both hemispheres, as measured using calcium imaging, voltage-sensitive dye imaging, and intrinsic signal imaging^16–18^. We therefore expected Robo3cKO mice to be severely impaired in our whisker side discrimination task. Surprisingly, Robo3cKO mice performed at levels comparable to their WT littermates (Figure 1). Moreover, performance depended specifically on activity in the hemisphere contralateral to a stimulated whisker, just as in WT (Figure 2). This too was surprising because Robo3cKO mice could in principle have solved the task based on evoked activity in either hemisphere. These observations could both be explained if the ectopic ipsilateral-whisker responses in wS1 were somehow filtered out by downstream areas.

In wS1, responses to ipsilateral and contralateral whiskers remained segregated at the level of individual units and linear probe penetrations (possibly columns) in cortex of Robo3cKO mice (Figure 3). This segregation facilitates a straightforward readout of individual single units (Figure 4) or of the population (Figure 5) to separate activity evoked by ipsi- or contralateral whisker stimulation, and thus to solve the task.

In contrast to wS1, whisker responses in wM1 of Robo3cKO mice were largely contralateral and similar to those of WT (Figures 6 and 7). This is consistent with ectopic ipsilateral responses in wS1 having been filtered out from the whisker processing stream at the level of wM1.

Our wM1 single-unit recording findings build on and contrast with prior work that used voltage-sensitive dye imaging and found bilateral evoked responses in wM1 of Robo3cKO mice^18^. Voltage-sensitive dye imaging can report subthreshold signals and synaptic inputs to a region, whereas our single-unit recordings only report spiking of local neurons. Together, these findings suggest that wM1 may receive both ipsi- and contralateral whisker input from wS1, but that only the contralateral inputs drive spiking. This could in principle arise from a plasticity mechanism that weakens the influence of ipsilateral-whisker synaptic inputs to the point that they fail to cause spiking in wM1.

We chose to examine wM1 because it is one of the two dominant corticocortical output pathways from wS1, along with whisker secondary somatosensory cortex (wS2)^10,11^. Both wM1 and wS2 have been implicated in whisker detection tasks^20–27^. In future work, it will be important to examine wS2 to determine what impact bilateral responses in wS1 have on wS2 activity and whether wS2 plays a role in compensating for the Robo3cKO mutation.

In addition to the lemniscal pathway running through VPM, information from the whiskers ascends via the paralemniscal pathway through the posterior medial thalamus (POm)^2,28–30^. This pathway likely remains unaffected in Robo3cKO mutant mice. POm sends inputs to wS1, wS2 and wM1^31–34^. Therefore, wM1 activity is shaped both by corticocortical projections from wS1 and by thalamocortical projections from POm. One hypothesis to explain the mostly normal behavioral task performance and wM1 response properties in Robo3cKO mice is that the paralemniscal pathway takes on a more dominant role vis-à-vis the lemniscal pathway. The relatively normal and contralateral responses in wM1 of Robo3cKO mice could arise from plasticity mechanisms that cause a relative strengthening of the influence of inputs from POm, at the expense of inputs from wS1. This hypothesis is not mutually exclusive with the hypothesis that ipsilateral responses are filtered out by selective weakening of inputs from ipsilaterally selective wS1 neurons.

Finally, our task relies on the processing of inputs from a single whisker on a given trial. The consequences of bilateral maps in wS1 could be different when processing simultaneous signals from multiple whiskers. In future work with Robo3cKO mice, it will be important to examine behavior and circuit function in the context of more complex, multi-whisker tasks.

## Acknowledgments

We thank Sara Stephenson for assistance with animal training. This work was supported by NIH grants R01NS084818 and 1U01NS115587-01. We thank Dr. Alain Chédotal (Institut de la Vision, INSERM, Sorbonne Université, Paris, France) for providing the initial breeding pairs of mice used in this study.

## Author contributions

V.B.C. and D.M.R. performed experiments. V.B.C. and Y-T.C., analyzed data. D.H.O. and R.S.E. planned the project. V.B.C. and D.H.O. wrote the paper with input from all authors.

---

**Figure S1:**
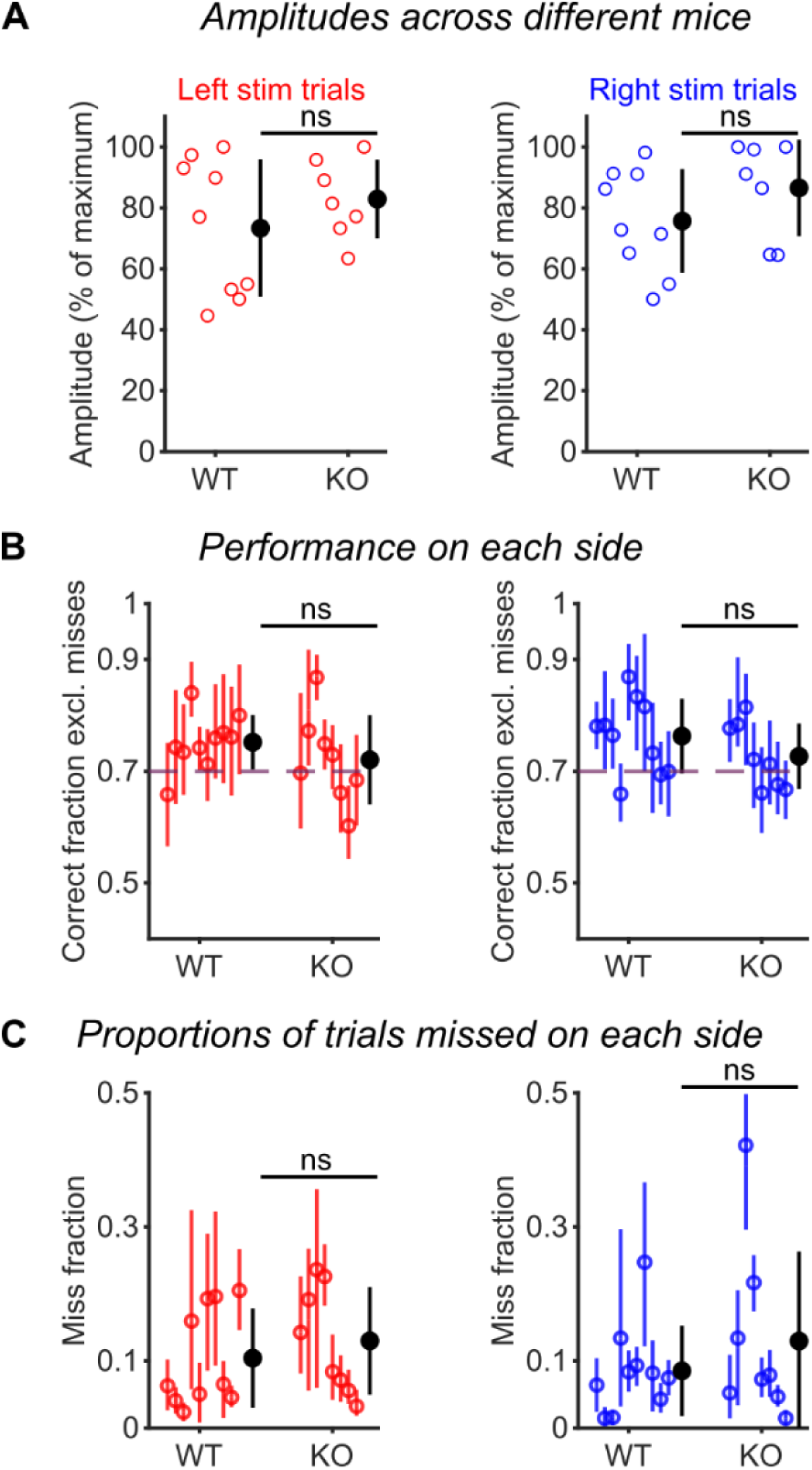
Similar task performance for Robo3cKO and WT mice. (A) Mean stimulus amplitudes for each mouse (open circles) were similar for both genotypes and for left (red) and right (blue) whiskers. Filled circles: mean ± SD across mice. Mann-Whitney U tests between genotypes show no significant (ns) differences. (B) Fraction of trials correct (excluding miss trials) was similar for left (red) or right (blue) whisker stimulation. Open circles: individual mice (mean ± 95% CI). Filled circles: mean ± SD across mice. Mann-Whitney U tests between genotypes show no significant (ns) differences. (C) Fraction of miss trials was similar for both genotypes and for left (red) and right (blue) whiskers. Conventions as in B.

**Figure S2:**
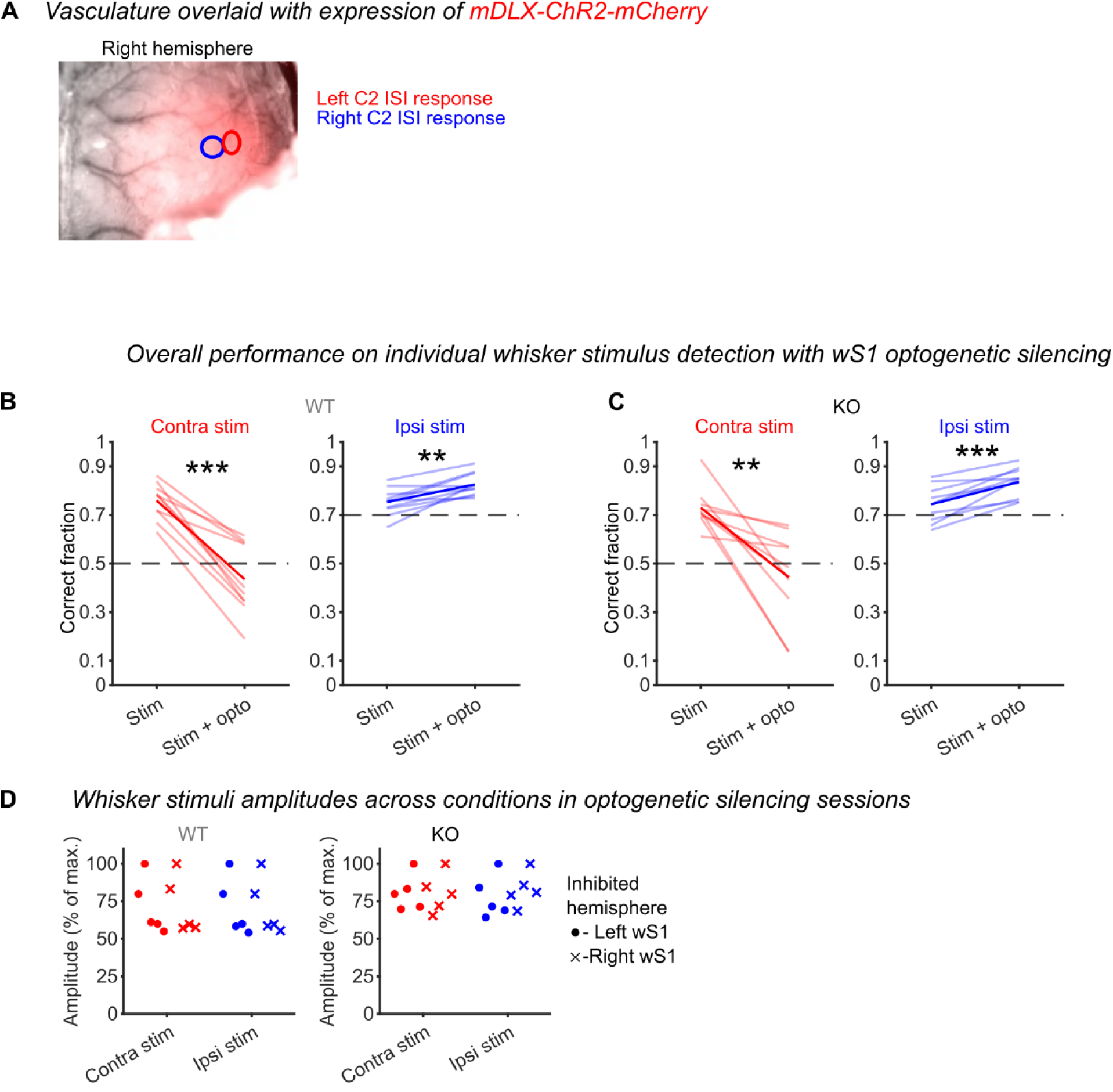
Robo3cKO and WT mice need contralateral S1 for whisker detection. (A) Example image from a Robo3cKO mouse showing channelrhodopsin-2 expression (red) with approximate locations of contralateral (red oval) and ipsilateral (blue oval) ISI responses indicated. (B) Performance for individual WT mice (thin lines) and the mean across mice (thick lines) for trials with and without unilateral optogenetic inhibition of wS1, and for stimulation of the C2 whisker contralateral (red) or ipsilateral (blue) to the inhibited hemisphere. (C) Same as panel B but for Robo3cKO mice. (D) Mean stimulus amplitudes were similar for both genotypes and for stimulation of the C2 whisker contralateral (red) or ipsilateral (blue) to the inhibited hemisphere. Plot symbols show individual mice but separately for inhibition of the left (circles) and right (crosses) wS1.

**Figure S3:**
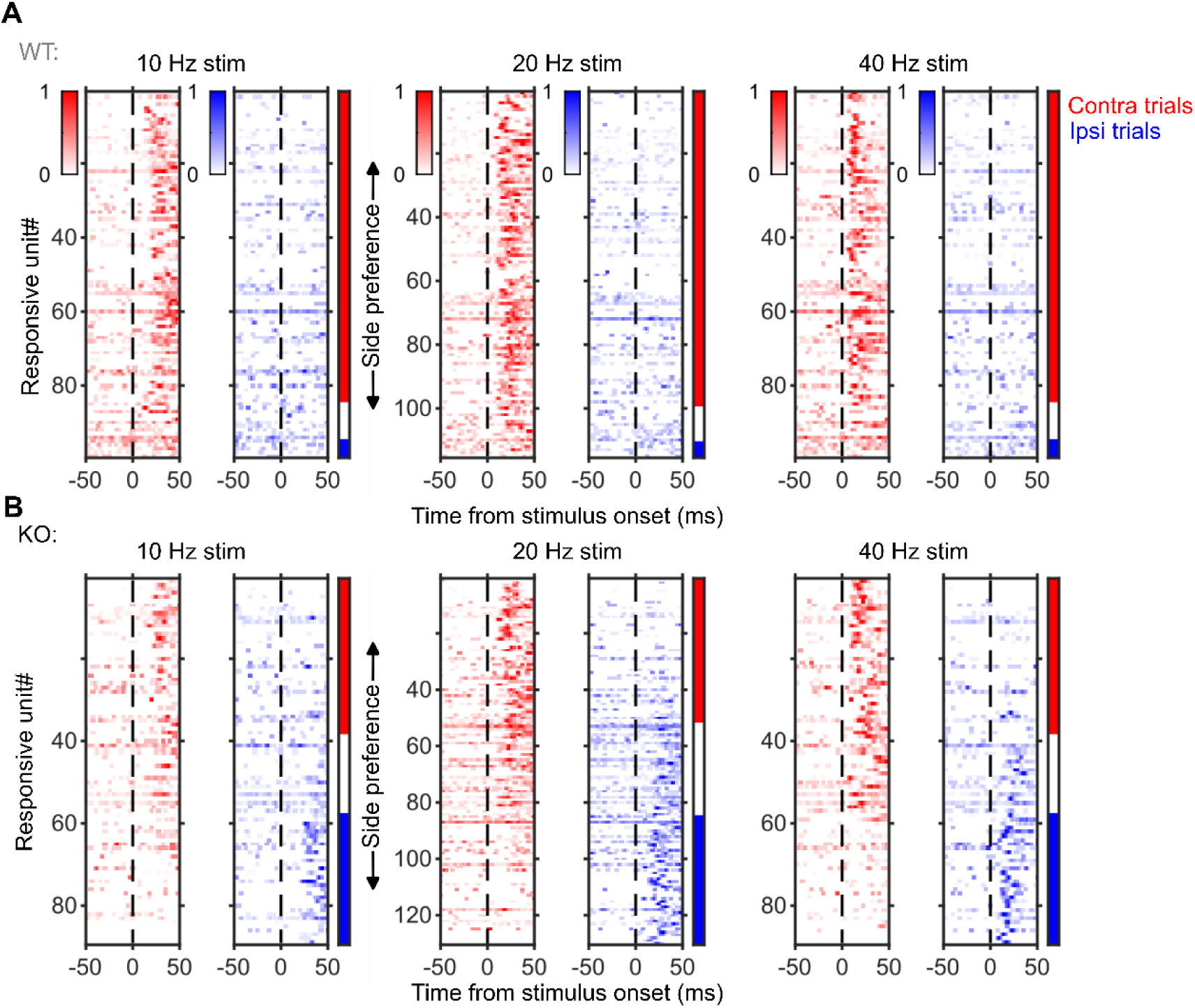
wS1 unit responses to stimulation at different frequencies in Robo3cKO and WT mice. (A) Similar to Figure 3C but showing results for three stimulation frequencies. Normalized response of wS1 units to contralateral (red) and ipsilateral (blue) stimuli of different frequencies (columns). Units are sorted by whisker preference based on laterality index values. Vertical bars at right indicate units with contralateral (red), bilateral (white) or ipsilateral (blue) preferences. (B) Same as A but for Robo3cKO mice.

**Figure S4:**
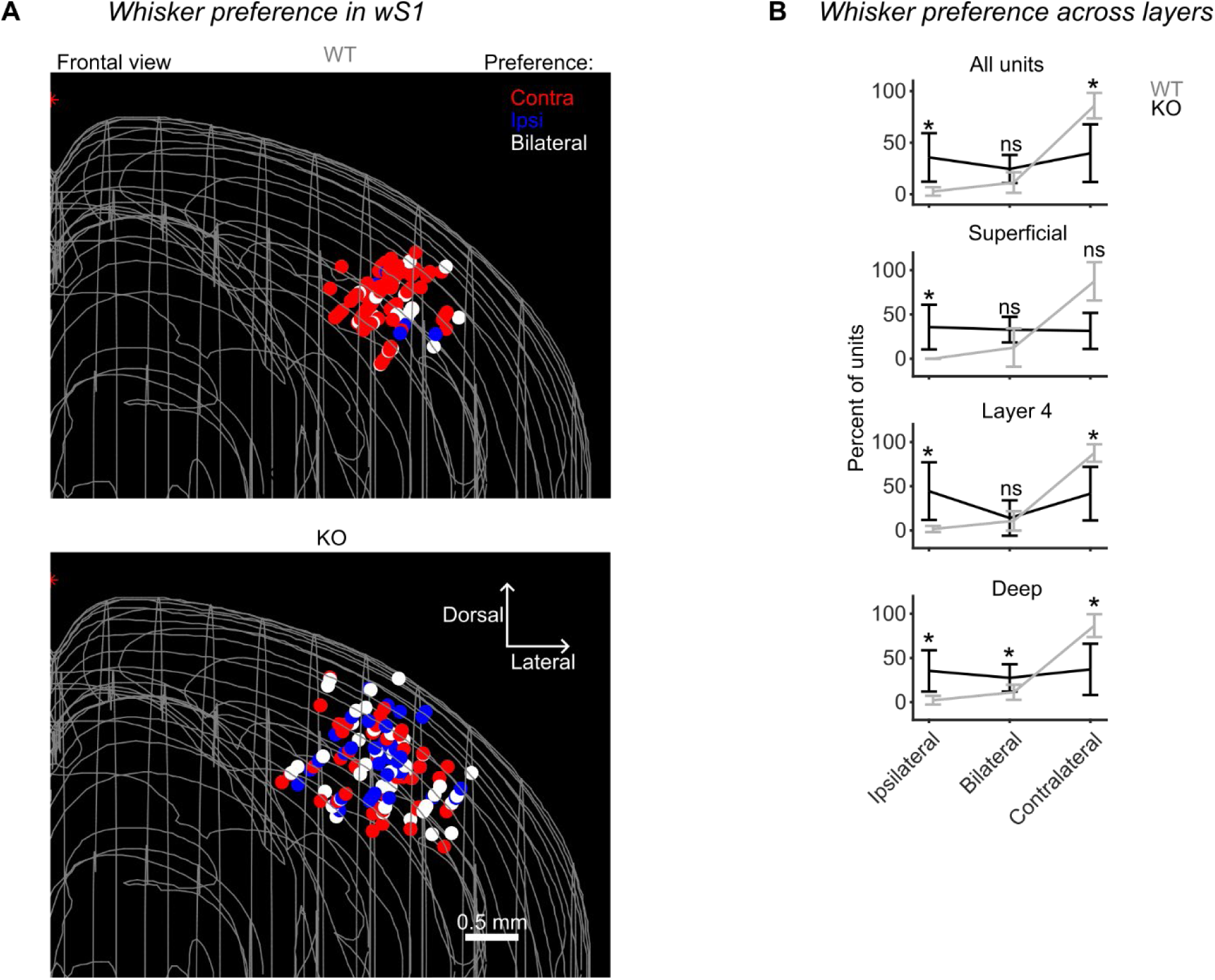
Locations of responsive units in wS1. (A) Depiction of locations of stimulus-responsive units in wS1 for WT (top) and Robo3cKO (bottom) mice registered using SHARP track^41^. Units are colored based on their preference for contralateral (red), ipsilateral (blue) or bilateral (white) C2 whisker stimulation. (B) Percent of wS1 units with each preference type found within nominal superficial, layer 4, or deep cortical layers (Methods). Mean ± SD across 6 mice for each genotype. Mann-Whitney U test between genotypes: *, p < 0.05; ns, not significant.

**Figure S5:**
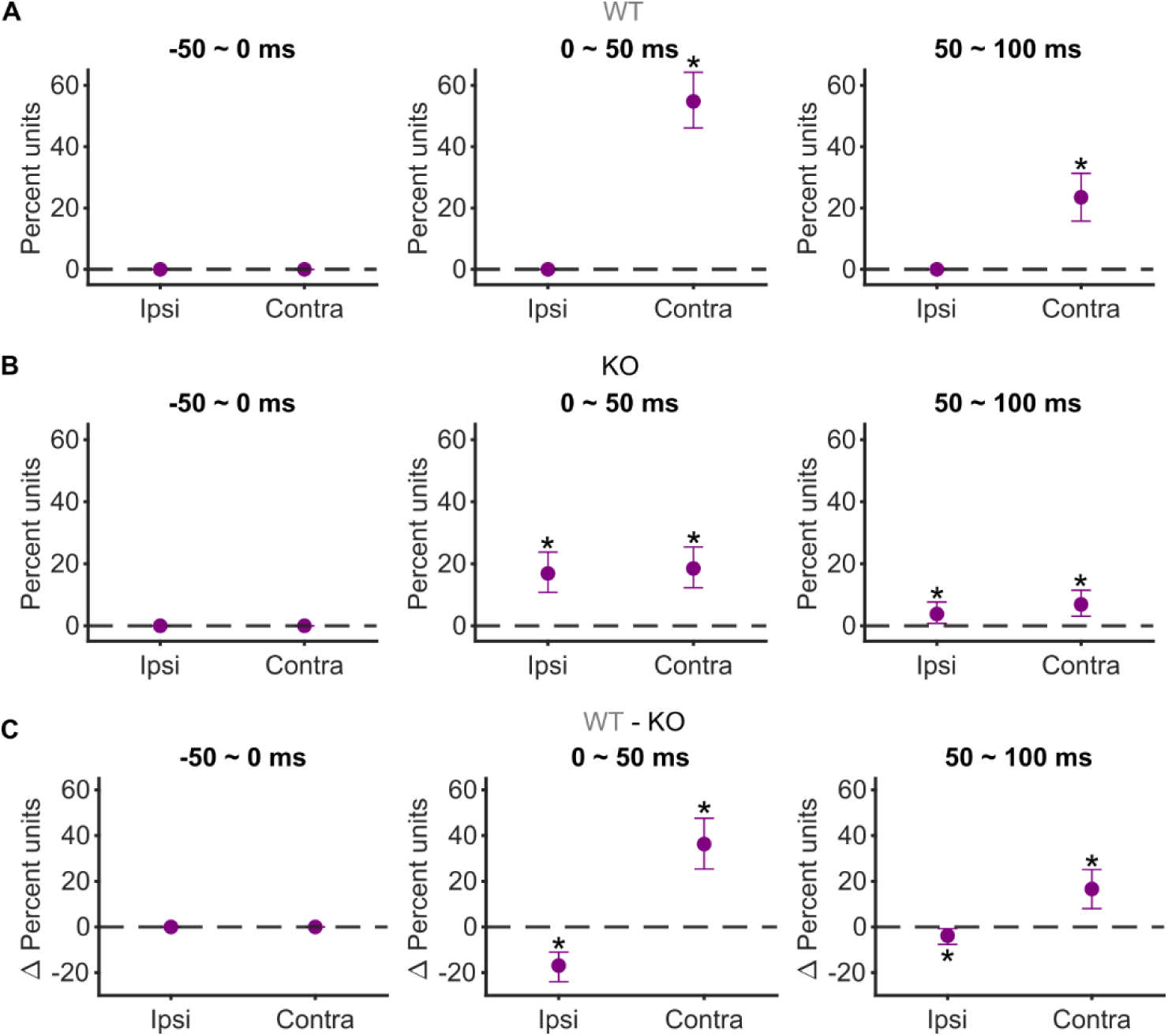
Confidence intervals for the percentage of units with whisker-side selectivity in wS1, related to. Figure 4A. (A) Percentages (± 95% bootstrap CI) of wS1 units with whisker-side selectivity calculated for different 50-ms time bins. Asterisks: confidence intervals that do not contain 0. (B) Similar to A but for Robo3cKO mice. (C) Similar to A but for the differences between WT and Robo3cKO mice.

**Figure S6:**
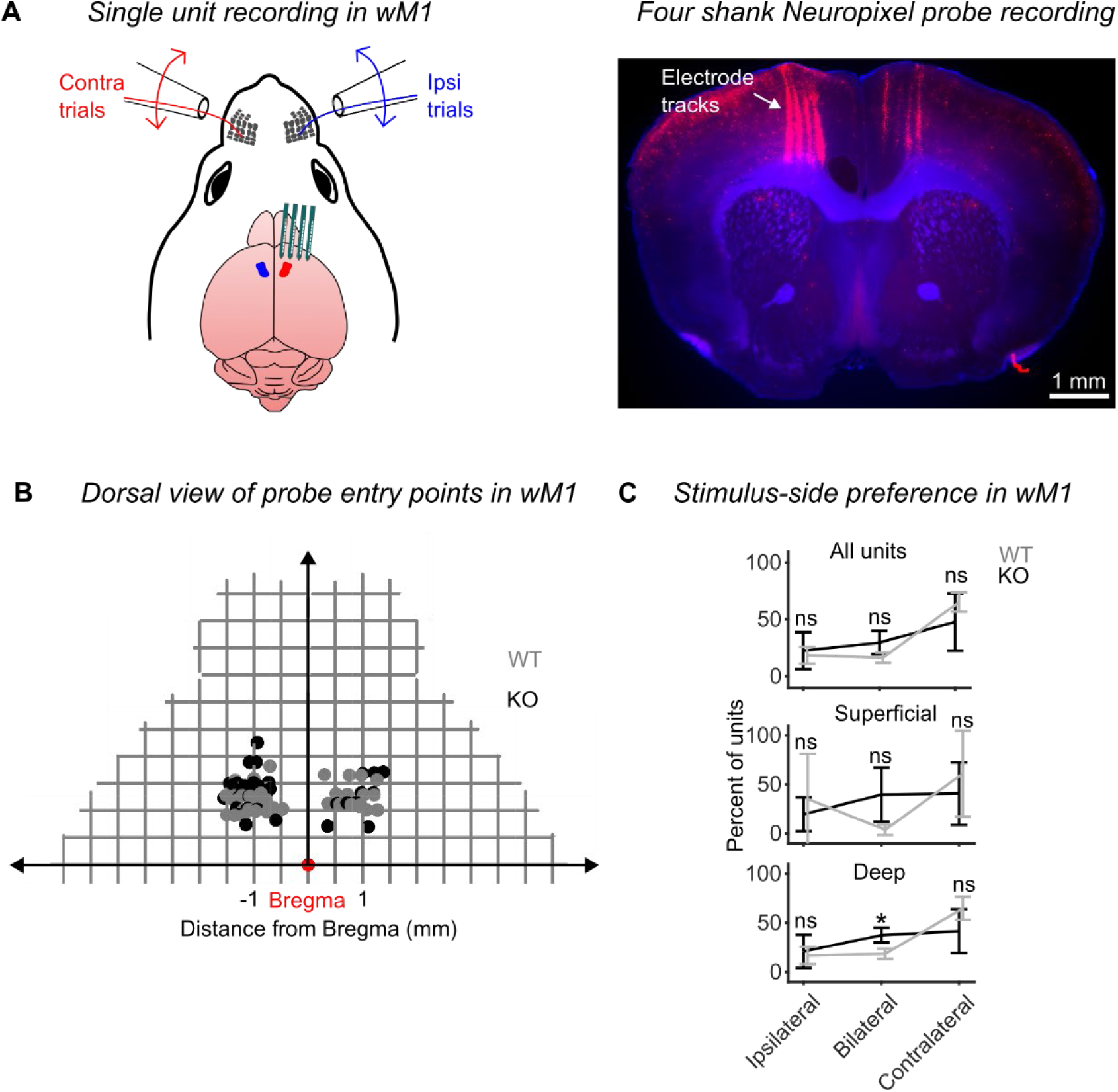
Locations of responsive units in wM1. (A) Single-unit recordings were performed using four-shank Neuropixel 2.0 probes targeted to motor cortex and coated with DiI (example tracts shown at right) to allow post hoc assignment of recording sites to wM1. (B) Dorsal view of probe penetration sites for all whisker responsive units in wM1 of WT (gray) and Robo3cKO (black) mice registered to the Allen Institute’s 2017 release of mouse atlas using SHARP track package^41^. (C) Percent of wM1 units with each preference type found within all units, nominal superficial or deep cortical layers (Methods). Mean ± SD across WT (gray, n = 3) and Robo3cKO (black, n = 4) mice. The contralaterally selective fraction was similar for WT and Robo3cKO mice. Mann-Whitney U test between genotypes: *, p < 0.05; ns, not significant.

**Figure S7:**
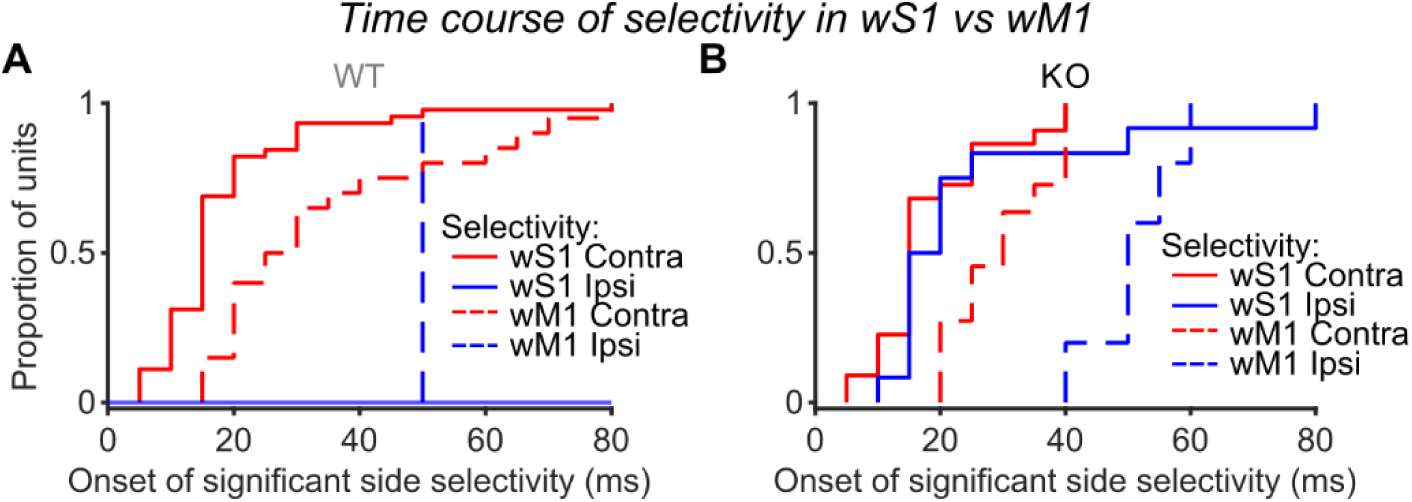
Onsets of whisker side selectivity in wS1 vs wM1. (A) Times relative to stimulus onset at which each unit from wS1 or wM1 of WT mice first showed a significant preference for the contralateral or ipsilateral C2 whisker. The onset of contralateral selectivity in wM1 is delayed compared to wS1. One-sided Kolmogorov-Smirnov test, S1 contra vs M1 contra: p < 0.0005. (B) Same as A but for Robo3cKo mice. The onset of ipsilateral selectivity occurs as early as that of contralateral selectivity in wS1 of Robo3cKO mice, and both occur earlier than contralateral selectivity in wM1. Two-sided Mann Whitney U test, S1 contra vs S1 ipsi: p = 0.27316. One-sided Mann Whitney U test, S1 contra vs M1 contra: p < 0.001, S1 ipsi vs M1 contra: p < 0.05.

**Figure S8:**
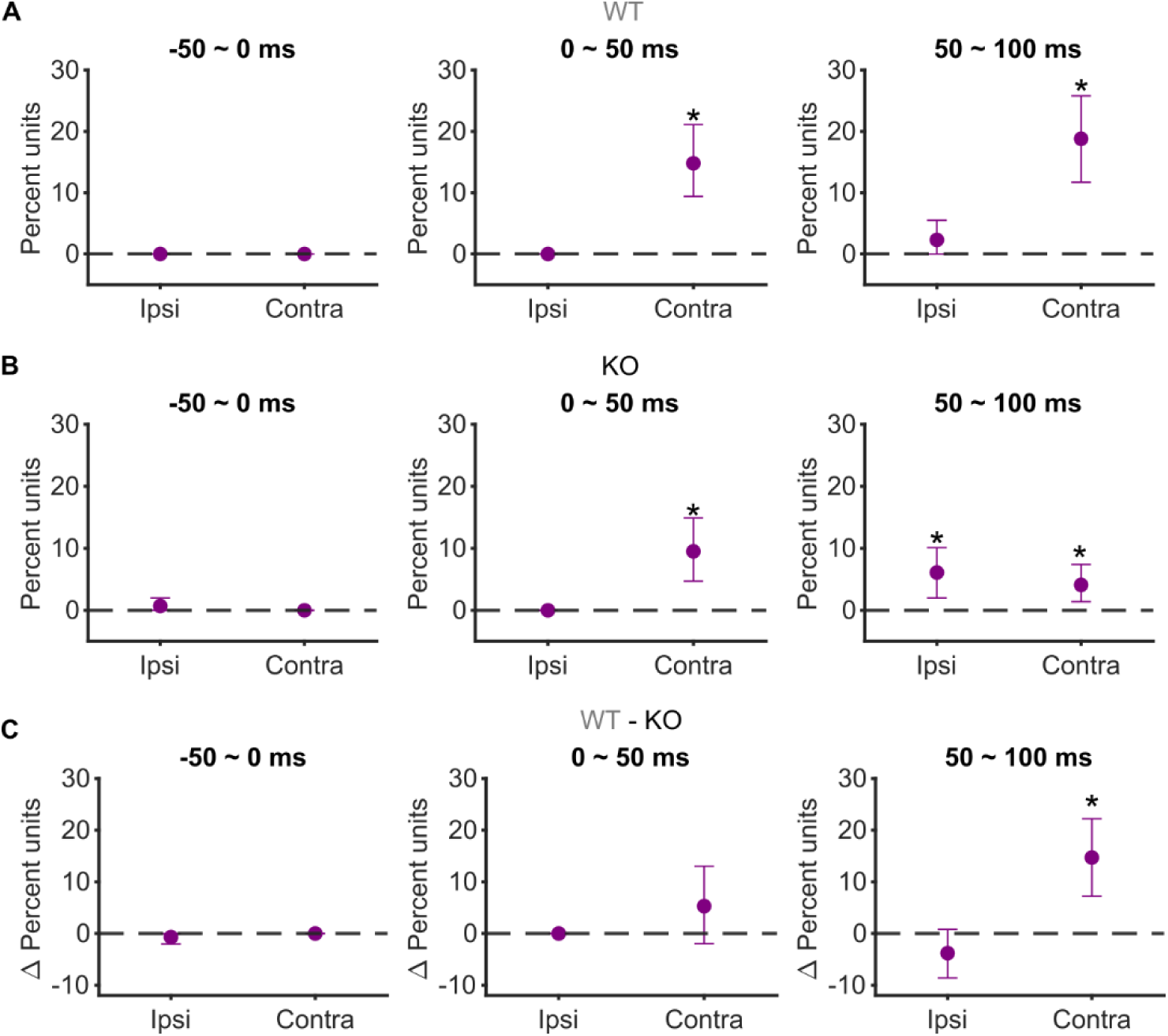
Confidence intervals for the percentage of units with whisker-side selectivity in wM1, related to. Figure 6C. (A) Percentages (± 95% bootstrap CI) of wM1 units with whisker-side selectivity calculated for different 50-ms time bins. Asterisks: confidence intervals that do not contain 0. (B) Similar to A but for Robo3cKO mice. (C) Similar to A but for the differences between WT and Robo3cKO mice.

## Methods

### Mice

All procedures were performed in accordance with protocols approved by the Johns Hopkins University Animal Care and Use Committee (protocols: MO21M195, MO24M185). Robo3cKO mice (Krox20:Cre;Robo3^lox/lox^) for experiments were generated by crossing Robo3cKO mice with Robo3^lox/lox^ mice as described previously ^7,35^. Cre-negative littermates were considered Robo3-WT (Robo3^lox/lox^, WT). Cre-recombination was confirmed by genotyping by following primers: 5’-AGC CCT TCA AGT ACC AGG GCC TGA C-3’, 5’-AGC GGT CCA GCA GGT ACA GCA TCA C-3’, 5’-ACC TGA TGG ACA TGT TCA GGG ATC G-3’and 5’-TCC GGT TAT TCA ACT TGC ACC ATG C-3’. Mice ranged in age 7-60 weeks at the time of experiments. Mice were housed in a vivarium with a reverse light-dark cycle (12 h for each phase) and were singly housed following surgical procedures and during behavioral experiments.

### Neonatal viral injections

All mice that underwent optogenetic inhibition experiments were given viral injections in wS1 as newborns (postnatal days 0-2) as previously described ^36,37^. Pups were cryo-anesthetized by placing them between wet towels on ice for 4-5 minutes ^38^. Adequate anesthetic depth was monitored by lack of movement and gentle paw pinch. A glass pipette with sharp tip (30-50 microns, typically beveled) was loaded with 250 nL of virus suspension (AAV1-mDlx-ChR2-mCherry-Fishell-3, 83898-AAV1), using a programmable automatic injector (Drummond Nanoject Ill). Anesthetized pups were restricted gently under a soft rubber strip secured by tape on a cold pack. Anatomical landmarks visible through the translucent skin were used to locate the center of wS1. Under a dissection microscope, the glass pipette was slowly inserted through the skin and soft cranium into wS1. Viral solution (∼200 nL) was slowly (∼50 nL/s) ejected from the pipette via positive pressure. Pups were kept on a warm pad until awake before returning to the home cage.

### Adult viral injections

Two mice (VC030114 and VC030211) in which neonatal injections failed to yield channelrhodopsin-2 expression in wS1 also underwent injections as adults, targeted using intrinsic signal imaging. Small craniotomies (300 µm diameter) exposed the area of wS1 responding to the C2 whisker in both hemispheres. Injections were done at 2 different depths from the pia (500 and 700 µm, 100 nL at each depth) to cover the whole depth of the cortex. The virus (AAV1-mDlx-ChR2-mCherry-Fishell-3) was injected slowly at ∼1nL/s.

### Surgery

Mice were prepared for head fixation and optogenetic silencing via clear-skull preparation^15^. Mice were anesthetized with isoflurane (1-2% in O_2_ at 1 liter/min, IsoSol), mounted in a stereotaxic apparatus (Kopf Instruments) with a thermal blanket (Harvard Apparatus), and administered Dexamethasone (2 mg/kg, subcutaneous). The scalp was cleaned after fur removal (Nair) using sterile saline and topical antiseptic (Betadine). Bupivacaine was injected under the scalp for local anesthesia. The dorsal skull was exposed by removing skin and periosteum. The temporal muscles were carefully detached from the lateral edges of the skull to allow access to the skull underneath. A thin layer of Metabond (C& B Metabond) was applied on the exposed skull surface and around the skin edges. A headpost was firmly cemented onto the posterior section of the skull using a thick layer of Metabond. A thin layer of cyanoacrylate glue (Krazy Glue) was applied over the Metabond to make it transparent. The clear skull surface was protected using a silicone elastomer (Body Double).

### Intrinsic signal imaging

Intrinsic signal imaging was performed to localize barrel columns corresponding to each spared C2 whisker after recovery from headpost surgery (∼7 days) and water restriction (7-10 days). Mice were maintained under light anesthesia with isoflurane (0.5–1%) and chlorprothixene (∼0.36 mg/kg; Sigma C1671, intramuscular). Intrinsic signal imaging was performed through the skull as described^39^. For each genotype, either left or right C2 whisker was stimulated using a glass pipette attached to a piezo, and responses imaged in each hemisphere. The rest of the whiskers were trimmed on each side of the face. Coordinates for the center of the response were aligned with an image of the vasculature to target optogenetic silencing experiments and electrophysiological recordings.

### Whisker side discrimination task

All behavioral experiments were conducted with head-fixed mice during the dark phase. The behavioral apparatus was controlled by BControl software (C. Brody, Princeton University). Seven to 10 days after surgery and 7-10 days before behavioral training, mice were allowed 1 ml of water daily. Body weight was maintained between 70-85% of the initial weight before the start of water restriction. On training days, mice performed the behavioral task until they reached satiety or for ∼1 hour, whichever occurred first. Mice consuming less than 0.7 mL of water were given supplemental water.

On the first day of training, mice were habituated to head fixation and the behavioral apparatus while being given free access to water via two lick ports positioned in front of their snouts and on either side of their midline. On all subsequent training days, a single whisker (always the C2 whisker) was threaded into a glass pipette loaded onto a piezo actuator (Q220-A4BR-1305YB, Piezo.com), which was driven by a piezo controller (MOT693A, Thorlabs). The glass pipette tip was approximately 2.5 mm from the follicle. All whiskers except the C2 whisker were trimmed near the base.

In the first two sessions, mice were trained to detect a stimulus (1 s sinusoidal deflections at 40 Hz, mean angular velocity of ∼1250 deg/s) on only one of the C2 whiskers (either left or right). The mice had to lick the associated lick port in a 1-s response window immediately following stimulus onset to get a water reward (∼6 μL). Licks that occurred within the first 0.2 s after stimulus onset (“grace period”) were ignored. To aid association of the correct lick port with the side of the whisker stimulus, the first session was generally an “auto-reward” session in which mice were given a reward at the end of the response window if they did not lick the incorrect lick port. An auditory white noise mask (cut off at 40 kHz; 80 dB SPL) was played continuously. The second session generally was without auto-rewards and the response window was extended to last until 2 s following stimulus onset (still with the 0.2 s grace period). In the subsequent two sessions, mice were trained with the whisker stimulus always on the other side. Following these four unilateral whisker stimulus sessions, mice were trained on bilateral sessions in which either the left or the right whisker was randomly chosen for stimulation. In the first two bilateral sessions, mice were stimulated with 40 Hz sinusoidal deflections for 1 s with auto-reward given at 0.3-2 s after stimulus onset. Auto-reward aid was removed starting with the third bilateral session. In the following sessions, stimulus strength was decreased by reducing the stimulus frequency to 20 Hz and the duration of sinusoidal deflection to 150 ms. The water reward was also gradually decreased to ∼3 μL to increase the number of trials performed.

Performance was assessed as the fraction of trials with correct responses after excluding trials when mice did not lick either of the two lick ports (“miss trials”). Miss trials were quantified separately. When the mice attained >75% trials correct with maximum amplitude, the amplitudes of the sinusoidal deflections were gradually adjusted downward within and across the sessions such that mice maintained performance within ∼70-75%. The stimulus amplitudes were independently adjusted for the left and right C2 whiskers. After reaching this performance criterion, mice proceeded with test sessions. Ten WT mice and eight Robo3cKO mice performed the whisker discrimination task. Behavioral test sessions lasted until mice were sated or for ∼1 hour. To ensure we analyzed only trials with stable task engagement, we excluded trials at the start and end of the session in which mice licked only one lick port or did not lick at all.

### Optogenetic inhibition of wS1

Following training and testing on the WSD task, a subset of mice (5 from each genotype) was prepared for and underwent optogenetic inhibition experiments. A visual masking flash was delivered for the duration of every trial (sinusoidal waveform at 40 Hz from -900 ms to +2000 ms relative to stimulus onset with a 200 ms ramp down; this covered the period from auditory cue to end of the response window) via a 470 nm LED (7007-PB000-D, LEDdynamics) placed in front of the eyes. Mice were considered trained to ignore the masking flash and ready for optogenetic inhibition once performance again reached 70-75% for three consecutive days. For inhibition experiments, an optic fiber was placed over wS1 of one hemisphere and centered on the location of the C2 responses obtained with intrinsic signal imaging. Photostimulation was randomly delivered on 20% of the trials and comprised a 40 Hz sinusoidal waveform from -200 ms to +2000 ms relative to stimulus onset with a 200 ms ramp down. Average power at the tip of the optic fiber (200 µm diameter, 0.22 NA, TM200FL1B, Thorlabs) was ∼8 mW. The optic fiber was coupled with a 473 nm wavelength laser (MBL-III-473-100mW, Ultralasers) and intensity controlled by an acousto-optic modulator (MTS110-A3-VIS, Quanta Tech) At least five sessions of unilateral wS1 inhibition were performed for each hemisphere. The side of unilateral inhibition was alternated across sessions. To ensure we analyzed only trials with stable task engagement, we excluded trials at the start and end of the session in which mice licked only one lick port or did not lick at all on the control (no-inhibition) trials.

Change in fraction correct performance due to optogenetic inhibition was calculated for each hemisphere after excluding miss trials. Confidence intervals were calculated using hierarchical bootstrapping (10000 iterations) with sessions and trials sampled randomly with replacement.

### Electrophysiology and response quantification

Mice that underwent optogenetic inhibition experiments were subsequently prepared for electrophysiology. Single-unit recordings were performed from mice trained on the WSD task, with the exception of two mice that were untrained (VC030401 and VC030402; see Supplemental Table 1). For silicon probe recording, a craniotomy was made over the recording site (∼0.6 mm in diameter for wS1 and 1x1 mm square for wM1). The skull was thinned along the perimeter of the craniotomy site using a spherical drill bit until it just broke through. The skull flap was carefully removed using a fine tungsten needle. Dura were carefully excised using a sterile tungsten needle (10130-05, Fine Science Tools) and absorbent points (#504, Henry Schein). The craniotomy was sealed and protected with a small layer of silicone elastomer (Kwik-Cast). A small craniotomy (∼0.3 mm in diameter) over the left hemisphere was made for implantation of a ground screw (4.2 mm anterior, 0.5 mm lateral to bregma). Craniotomies were made one day before electrophysiological recordings. If a mouse underwent electrophysiological recordings from wM1, these were obtained in both hemispheres before performing craniotomies for wS1 recordings. For recording sessions, mice were maintained under light isoflurane anesthesia (0.7% in O_2_ at 1 liter/min). Breathing rate was visually monitored to be ∼1 breath per second.

wM1 recordings were targeted stereotactically using coordinates relative to Bregma (AP: +0.8-2 mm; ML: 0.5-1.5 mm) using 384-channel Neuropixel 2.0 four-shank probes coated with CM-DiI (Invitrogen). With recording sites distributed across all four shanks we could acquire data across 700 µm of depth at a time, although the electrodes spanned ∼1400 um of depth. To cover the whole span of the cortical depth, we divided each session into two epochs of ∼300 trials. The first epoch consisted of recording from the bottom 700 µm of cortex for at least 300 trials, followed by a second epoch covering the most superficial 700 µm of the cortex. From each hemisphere, 3-4 recording penetrations were made to completely span the area around wM1.

wS1 recordings were targeted using intrinsic signal imaging coordinates. Linear 64-channel probes (H3, Cambridge NeuroTech) or 384-channel Neuropixel 2.0 single-shank probes were coated with CM-DiI (Invitrogen) or DiD (5-10 mg/mL, Invitrogen). Probe arrays were inserted into the cortex at ∼30-35 degrees from the vertical.

After probe insertion, the brain was covered with a layer of 2% agarose and ACSF and left for ∼15 minutes prior to recording. Whisker stimuli were delivered using the same apparatus used during the WSD task and alternated between the left and right C2 whiskers. On each trial the stimulus was randomly chosen from a set comprising 1, 2 or 3 deflections with a 10, 20, or 40 Hz sinusoidal waveform. Recordings of wS1 units in two mice (VC030107 and VC030206; see Supplemental Table 1) were done in response to only 20 Hz stimuli.

Neural signals and behavioral timestamps were recorded with an Intan system (RHD2000; Intan Technologies) for 64-channel H3 probe recordings or with SpikeGLX (Bill Karsh, HHMI Janelia) for Neuropixel 2.0 probe recordings. Neural signals were sampled at 30 kHz. Kilosort 2.5^40^ was used for spike sorting with manual curation using Phy (github.com/cortex-lab/phygithub.com/cortex-lab/phy). Units were excluded from further analysis if the rate of inter-spike-interval violations exceeded 0.5% in a 2-ms window, the false positive rate exceeded 15%, or cut-off of the Gaussian distribution of amplitudes in a cluster exceeded 10%. These quality metrics were calculated using ecephys (https://github.com/jenniferColonell/ecephys_spike_sorting).

Neural spike rates were calculated in 2.5-ms bins and smoothed with a Gaussian kernel (5 ms). Units that were not significantly responsive to either C2 whisker stimulus were excluded from analyses.

For heat map plots (e.g. Figure 3C), the spike rate of each unit was normalized to the maximum of its mean responses across both whisker stimulus sides.

Laterality index (LI) values for each unit were calculated as LI = (contra response – ipsi response) / (contra response + ipsi response). The contra response was the absolute difference between spike counts calculated over 150-ms windows immediately after and before stimulus onset across all contralateral stimulus trials. The ipsi response was calculated similarly for ipsilateral stimulus trials.

### Histology

Mice were perfused transcardially with PBS followed by 4% PFA in 0.1 M PB. The brain was fixed in 4% PFA overnight. The brain was then embedded in 5% agarose in PBS. A vibratome (TedPella EasiSlicer) was used to cut coronal sections of ∼90 or ∼120 μm thickness. Sections were mounted and imaged on a fluorescence microscope (Zeiss AxioZoom V16). Images with DiI and DiD fluorescence were acquired to reconstruct recording locations.

### Unit location estimation

To estimate the locations of recorded neurons, coronal brain sections containing electrode tracts were registered to the Allen Brain Atlas (10 µm voxel 2017 release) using the SHARP track package^41^. The registered location of the electrode array was determined for each recording site, and the channel with the highest waveform power was used as the estimated coordinate of the unit. For analysis of whisker preferences vs cortical depth in wS1 (Figure S4), we assigned units to nominal “superficial” (0-518 μm), “layer 4” (418-588 μm) or “deep” (588-1154 μm) layers^34,42^. For wM1 (Figure S6), we assigned units to nominal “superficial” (≤ 285 μm) or “deep” (>285 μm) layers^43^.

### Single-unit whisker discriminability analysis

ROC analysis was used to calculate how well an ideal observer could use the trial-by-trial activity of a single neuron to discriminate between trials with contralateral vs ipsilateral whisker stimulation. Units that were responsive to the 20 Hz, 150-ms duration stimulus were included in this analysis. AUC values were calculated (MATLAB “perfcurve”) in 50-ms time bins (Figure 4A and Figure 6C). For each time bin, a Bonferroni corrected 95% CI for the AUC was obtained by bootstrap (1000 iterations). Units were considered significantly discriminative when this CI did not include the chance level (= 0.5). To determine the onset times of discriminability (Figure 4B-C, Figure 6D-E and Figure 6), we used a 5-ms bin and defined the onset as the time of the first bin in three consecutive significant bins.

### Unit categorization

Units were “responsive” to a particular stimulus if a two-tailed Wilcoxon signed-rank test showed a significant (p < 0.01) difference between spike rates calculated over 150-ms windows immediately before and after stimulus onset.

Unit “preference” for the side of whisker stimulation was determined based on LI values:

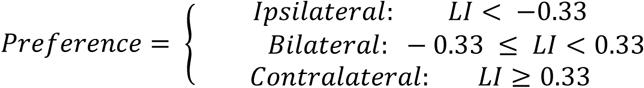

A unit was “selective” for the contralateral C2 when its AUC value was significantly > 0.5, and for the ipsilateral C2 when its AUC value was significantly < 0.5.

### Population decoding analysis

We used linear discriminant analysis (LDA; MATLAB “fitcdiscr”) to measure how well population activity from simultaneously recorded neurons could be used to decode the side of a whisker stimulus (contralateral vs ipsilateral). Sessions with at least three responsive units were included in this analysis. We used 50-ms time bins for wS1 (Figure 5). Given the slower development of selectivity in wM1 compared with wS1, we used 100-ms time bins for wM1 (Figure 5H). Shuffled data were generated by shuffling the contralateral vs ipsilateral trial labels. Confidence intervals were calculated using a hierarchical bootstrap method. To prevent the same trials from appearing in both the training and test datasets after bootstrapping, we first performed a 10-fold cross-validation partition and then bootstrapped trials within the training and test datasets separately. For each bootstrapping iteration (100 iterations total), classification accuracy was averaged across sessions.

### Statistics

We report data as the mean ± 95% confidence interval except where noted. Statistical tests were two-tailed unless otherwise noted. Confidence intervals specified as “hierarchical” were calculated using a nonparametric hierarchical bootstrap method (1000 iterations) that simulated the data generation process by incorporating variability at different levels, including mice, sessions, neurons, and trial types as applicable. Binomial tests in Figure 2 were performed (MATLAB “binopdf”) across mice on observations pooled from left and right wS1.

